# Family reunion via error correction: An efficient analysis of duplex sequencing data

**DOI:** 10.1101/469106

**Authors:** Nicholas Stoler, Barbara Arbeithuber, Gundula Povysil, Monika Heinzl, Renato Salazar, Kateryna Makova, Irene Tiemann-Boege, Anton Nekrutenko

**Affiliations:** Graduate Program in Bioinformatics and Genomics, The Huck Institutes for Life Sciences, The Pennsylvania State University, University Park, PA, USA; Department of Biology, The Pennsylvania State University, University Park, PA, USA; Institut für Biophysik, Johannes Kepler Universität, Linz, Österreich, (Austria)

## Abstract

Duplex sequencing is the most accurate approach for identification of sequence variants present at very low frequencies. Its power comes from pooling together multiple descendants of both strands of original DNA molecules, which allows distinguishing true nucleotide substitutions from PCR amplification and sequencing artifacts. This strategy comes at a cost—sequencing the same molecule multiple times increases dynamic range but significantly diminishes coverage, making whole genome duplex sequencing prohibitively expensive. Furthermore, every duplex experiment produces a substantial proportion of singleton reads that cannot be used in the analysis and are, technically, thrown away. In this paper we demonstrate that a significant fraction of these reads contains PCR or sequencing errors within duplex tags. Correction of such errors allows “reuniting” these reads with their respective families increasing the output of the method and making it more cost effective. Additionally, we combine error correction strategy with a number of algorithmic improvements in a new version of the duplex analysis software, Du Novo 2.0, readily available through Galaxy, Bioconda, and as the source code.

## Introduction

Numerous, often clinically important, research scenarios require detection of sequence variants that are present in a minute fraction (10^−5^−10^−9^) of molecules under study. Examples include detection of cancer-related mutations in liquid biopsies, identification of fetal DNA in a mother’s bloodstream, assessing dynamics of the immune system, tracing mutational landscape of bacteria through the evolution of antibiotic resistance, studying genomic changes in viral pathogens and many others (for a comprehensive review see (Salk et al. 2018)). Conventional approaches, where a sample is sequenced and resulting reads are aligned against a reference genome to find differences, are ill suited for variants present at frequencies below 1% (Rebolledo Jaramillo et al. 2014; Schmitt et al. 2015). A number of techniques has been developed to circumvent this issue with Duplex Sequencing (DS) being currently the most sensitive (Schmitt et al. 2012; Salk et al. 2018). DS is based on using unique tags (also called barcodes throughout this manuscript) to label individual molecules of the input DNA. During amplification steps that are required for the preparation of Illumina sequencing libraries, each of these molecules gives rise to multiple descendants. The descendants of each original DNA fragment are identified and grouped together using tags—one simply sorts tags in sequencing reads lexicographically and all reads containing the same tag are bundled into a *family*. These families (usually with three members or more) form single stranded consensus sequences (SSCS) for the forward or the reverse strand, respectively. Complementary SSCSs are then grouped to produce duplex consensus sequences (DCSs; see Fig. 1). A legitimate sequence variant is found in the majority of the reads within a family. In contrast, sequencing and amplification errors will manifest themselves as “polymorphisms” within a family and so can be identified and removed (yellow rectangles in Fig. 1).

**Figure 1.**
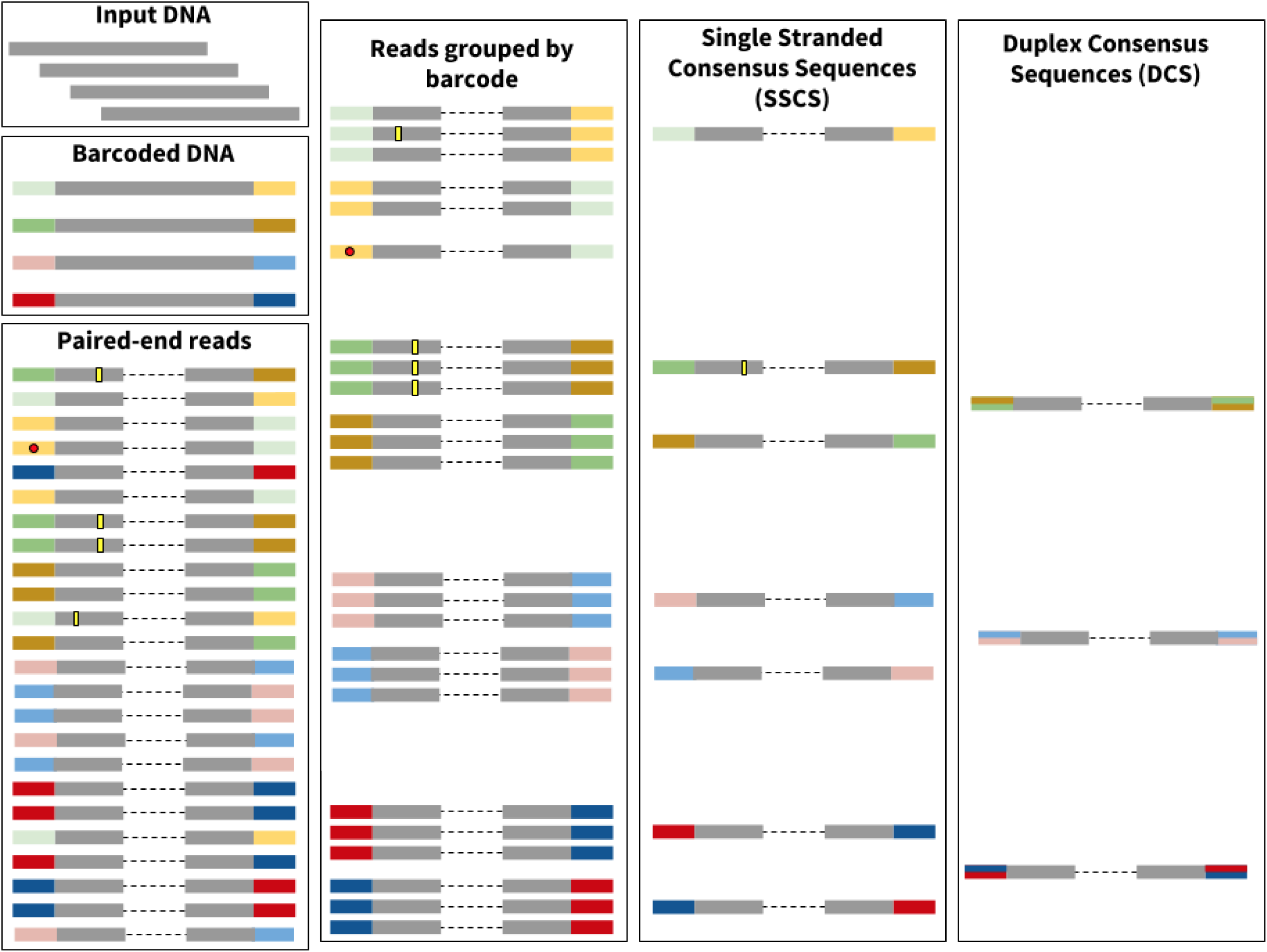
Effect of errors on the Duplex Sequencing procedure. Here input DNA is sheared and barcodes are ligated to the ends of the DNA molecules (colored rectangles in **Barcoded DNA**). After paired-end library preparation and sequencing each original molecules gives rise to multiple reads (**Paired-end reads** pane). This process also inadvertently generates sequencing errors represented by yellow rectangles and red circles. The yellow rectangles and red circles are used to depict errors arising inside read compartments corresponding to original DNA and adapters, respectively. Reads are then grouped by barcode to produce “families”. In this example each family is required to contain at least three reads. As shown here one of the reads contains an error (red circle) within the barcode. The error makes this particular barcode different from others. As a result it cannot be added into the family and remains a singleton (the error correction algorithm described here was developed specifically to correct such errors and allow singletons to be joined with their respective families). Each family is subsequently reduced into a Single Strand Consensus Sequence (SSCS) and each respective SSCS is merged with its counterpart from the opposite strand to generate Duplex Consensus Sequences (DCS).

Despite its power DS is a complex technique. Reliable identification of sequence variants requires each initial fragment to form a family with at least three members for each strand. To achieve this, it is necessary to precisely quantify the amount of input DNA during the library preparation step. Too much DNA results in small family sizes and makes variant identification impossible, while too little creates very large families at the expense of sequencing coverage. Furthermore, because DS barcodes are a part of the sequencing read, they accumulate PCR and sequencing errors. These errors prevent matching barcodes and therefore artificially split DS read families (red dots in Fig. 1) decreasing the efficiency of the procedure. In this manuscript we describe a new, efficient approach to the analysis of DS data that includes barcode error correction. It significantly improves the yield and performance of the technique. We also describe new quality control approaches designed to increase output of DS experiments.

## Results and Discussion

### Datasets

To test our results we used two previously published datasets. The first dataset was produced by Schmitt et al. (Schmitt et al. 2015), who employed DS to identify a rare mutation at the *ABL_1_* locus responsible for resistance to a chronic myeloid leukemia therapeutic compound imatinib. The second dataset was produced by our group as a part of an experimental evolution study where DS was used to track frequencies of adaptive mutations in plasmid _p_BR_322_ (Mei et al. 2018).

### Barcode errors result in lost reads

Typical DS tags are randomized 12-mers. Since each DNA fragment is labeled by two tags, one at each end, there are theoretically 4^(12+12)^ unique combinations. However, the input DNA in a standard DS experiment contains ~10^6^ – 10^11^ molecules creating a large tag-to-input excess (4^24^ ≫ 10^11^). Because of such excess it is, theoretically, highly unlikely to observe distinct input DNA molecules tagged by barcodes that are highly similar to each other.

To confirm this expectation we have selected 1,000 random duplex tags from the two datasets discussed above and compared them against each other (Fig. 2A; 1,000 tags were selected to speed up the computation; see Methods). Because tags are used to group reads into families, the family size is known for each tag. We also know if tags form single strand families (SSCS) and which of these, in turn, form duplex families. By comparing each of 1,000 tags against themselves we can calculate the number of differences among all tags (the Hamming distance referred to as “HD” in the remainder of this manuscript). Most tags are different by 5-7 nucleotides, which is consistent with our expectation. However, a substantial fraction of tags (85 and 494 out of 1,000 for *ABL_1_* and _p_BR_322_ datasets, respectively; Fig. 2A) differed by just a single nucleotide, which, based on the large excess of unique tags to molecules mentioned above, is unexpected. A likely explanation to this is that tags differing by a single nucleotide are in fact derived from the same barcode and the single difference is the result of PCR or sequencing error. An observation further supporting this explanation is that almost a half of the reads containing these tags (grey areas of the bars in Fig. 2) are singletons—do not form families with any other reads. Reads containing tags forming larger families (with at least two members) must have occurred in an earlier step given that more than one read is affected, and are therefore likely PCR errors. The core issues with errors within tags is that they prevent combining reads into families as reads containing such tags cannot be grouped into their respective SSCS and are therefore lost reducing the overall efficiency of a duplex experiment (Fig. 1). Being able to correct these errors is thus critical to improving cost effectiveness of duplex sequencing.

**Figure 2.**
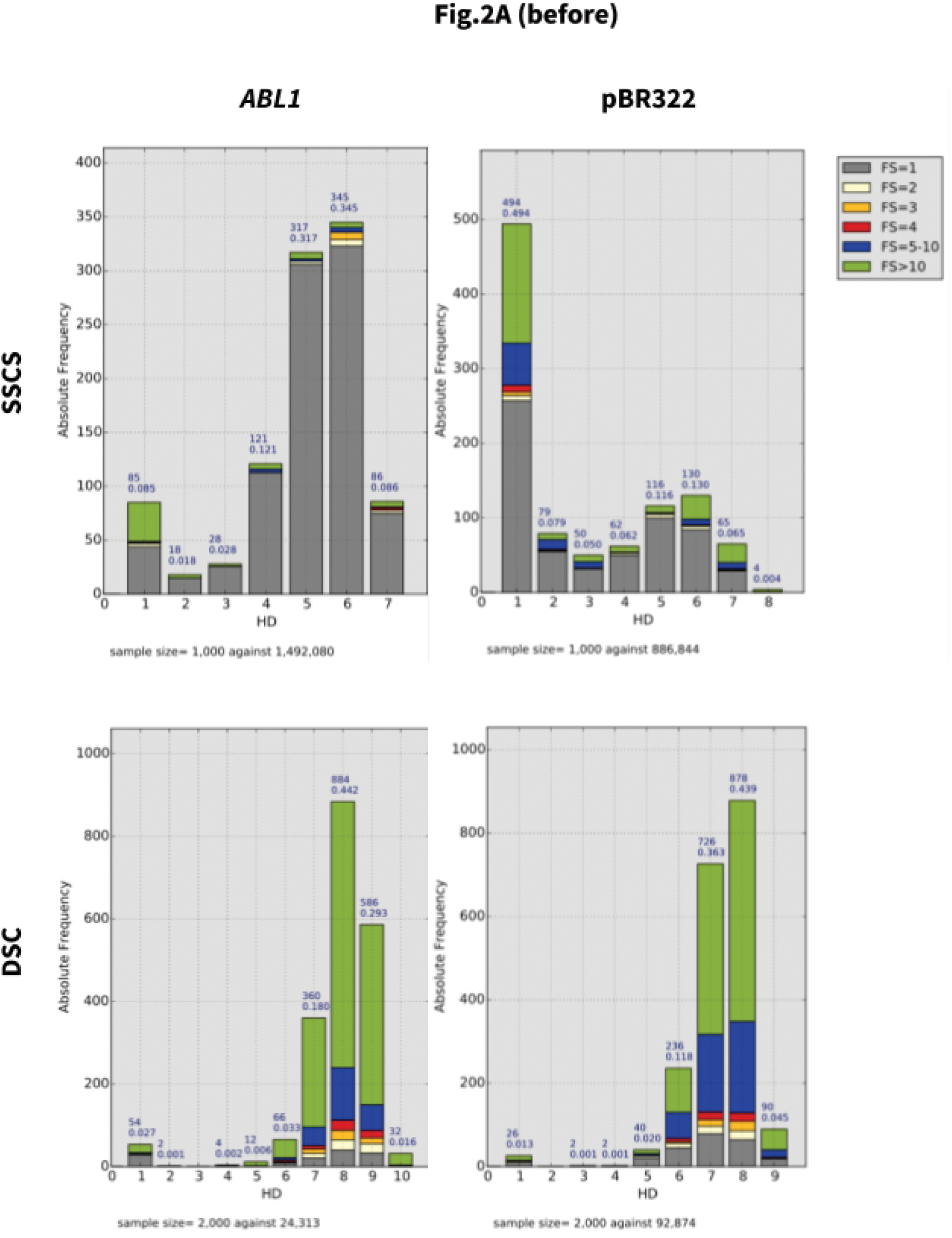
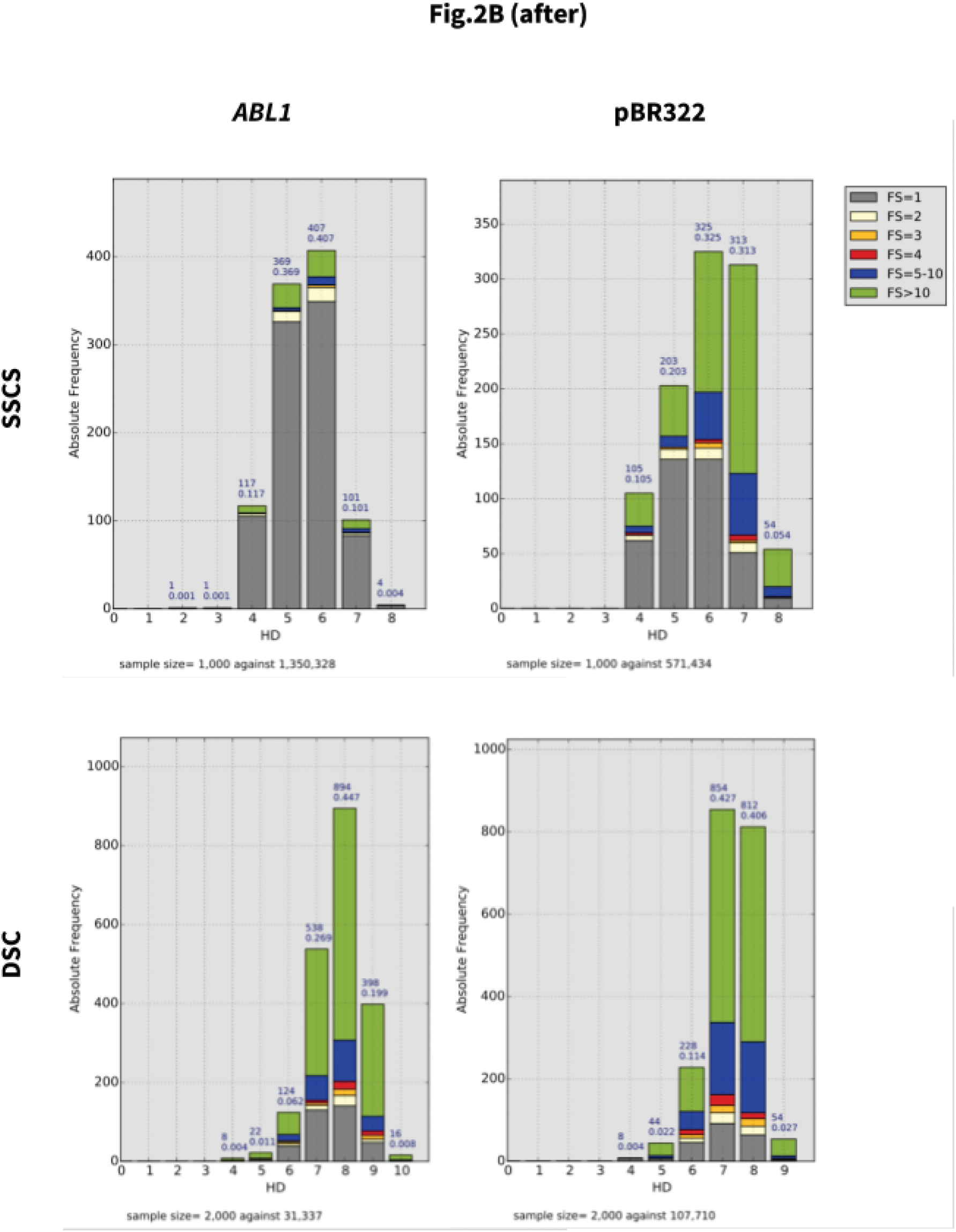
Error correction increases yield of Duplex Sequencing. In each figure top two graphs represent counts of SSCSs and bottom two show counts of DCSs. **(A).** Before error correction. **(B)**. After error correction.

### Barcode error correction increases yield

Forming families of reads descended from the same original fragment requires grouping reads by barcode (Fig. 1). This is straightforward when no sequencing errors are present and can be done by simple lexicographic sorting. Yet as we have shown in the previous section, errors are widespread and this eliminates sorting as a legitimate analysis strategy. An alternative approach will involve performing all-versus-all comparison of all barcodes to identify those that differ by one or two nucleotides and further checking them to see if they are potentially derived from the same DNA fragment with differences being introduced by PCR or sequencing errors. The challenge is that the all-versus-all comparison has *O*(*n*^2^) time complexity and thus is prohibitive as a routine analysis strategy. There are several tools that approach this problem in different ways. The most common strategy is to reduce the search space by first aligning the raw reads to the reference genome. One can then consider barcodes of only those reads that align to one region of the reference. This does not change the time complexity of the search, but reduces the search space from the millions of barcodes in the entire sample to the dozens that may be aligned to a particular genomic location. Several tools are available which use this strategy (Smith et al. 2017; Xu et al. 2018; Fennell and Homer 2018). However, reference-based approaches are inevitably biased and it was our main impetus to avoid the use of a reference sequence (Stoler et al. 2016). Alternatively, a strategy implemented in MAGERI, a tool which does not require a reference sequence to form consensus sequences, is able to perform efficient barcode error correction with the use of a custom seed-and-extend alignment algorithm (Shugay et al. 2017, 2014). However, it only forms single-strand consensus sequences, not the duplex consensus sequences required in our analysis.

To overcome these limitations we have adopted Burrows-Wheeler *k*-mer indexing implemented in Bowtie (Langmead et al. 2009) to quickly perform all-versus-all comparison of duplex tags. We are using the original version of Bowtie (not Bowtie2) that was optimized for very short reads. Specifically, we create an FM-index for all barcodes in the sample and then align individual barcodes (as if they were reads) against that index. Results of this alignment are represented as a graph where each vertex corresponds to a barcode. An edge is drawn between two vertices if an alignment exists between two barcodes. An alignment should have mapping quality and edit distance above a user-defined threshold with default values set to 20 and 3, respectively (in the discussion below we vary edit distance values from 1 to 3). The resulting graph contains a large number of disconnected clusters, each of which theoretically represents a single barcode together with all its derivatives created due to PCR and sequencing errors. A correct barcode can therefore be chosen by picking the vertex with the highest number of reads associated with it. To assess effectiveness of this error correction strategy we have developed a tool for producing simulated DS data (see Methods). Using this simulator we produced 400,000 duplex reads and analyzed them using our error correction approach. We then proceeded to calculate how many families (and thus, DCSs) were added to the analysis because of the correction. This increase in yield—the most important consequence of error correction— was substantial. The 400,000 simulated duplex reads produced 43,344 DCSs without correction. Running error correction by setting edit distance to one, two, or three mismatches resulted in 52,896, 53,420, and 53,454 DCSs, respectively. This constituted a 23% increase in yield (at three mismatches) compared the an uncorrected analysis. Effectively the error correction algorithm “shrinks” the pool of singletons (family size, FS, of 1) by reuniting them with families containing correct barcodes, increasing the likelihood that a group of reads surpasses the minimal member number (family size [FS] ≥ 3) for calling a SSCS.

Next, we proceeded to test our approach on real duplex sequencing datasets we used above. We specifically explored if tag error correction improves the number of consensus bases in SSCS and DCS, when allowing for 1, 2, or 3 mismatches in the tags. The results of error correction are summarized in Table 1, Fig. 2, and Fig. S1. The error correction decreased the number of singletons (FS ≥ 3) while increasing the numbers of DCSs by re-incorporating singletons into duplex families. This was particularly striking in _p_BR_322_ dataset, where the number of DSC increased from 77,164 to 89,513. One can also see that increasing the edit distance during error correction to 2 or 3 did not have such a drastic effect in reducing the number of singletons and increasing the overall SSCS and DCS (Fig. S1).

**Table 1.** Effect of error correction on duplex datasets analysis

| Sample | ABL1 |  |  |  | pBR322 |  |  |  |
| --- | --- | --- | --- | --- | --- | --- | --- | --- |
| # errors | 0 | 1 | 2 | 3 | 0 | 1 | 2 | 3 |
| SSCS ab | 38,493 | 37,803 | 37,007 | 36,280 | 84,231 | 81,929 | 78,481 | 73,647 |
| SSCS ba | 38,202 | 37,496 | 36,772 | 36,080 | 84,085 | 81,741 | 78,234 | 73,160 |
| DSC | 20,745 | 21,299 | 22,151 | 23,180 | 77,164 | 80,640 | 84,359 | 89,513 |

### New alignment engine improves consensus generation

The first version of Du Novo had a number of limitations resulting in poor performance. It was taking close to 24 hours to analyze a single duplex experiment. There were two primary reasons for this: the use of MAFFT aligner and inadequate parallelization strategy for executing multiple consensus generating jobs.

First, we sought to increase the performance of consensus generation step by employing a different multiple alignment tool that can be integrated into Du Novo codebase. We evaluated two candidate tools: SeqAn (https://github.com/seqan/seqan) and Kalign2 (Lassmann et al. 2009). SeqAn is a library of algorithms, including a multiple sequence aligner, specifically written to be incorporated into other genomics tools. Written in C++, it can be compiled and its functions called from Python. Kalign2 is an aligner using the Wu-Manber approximate string-matching algorithm (Wu and Manber 1992) to significantly speed up alignment while maintaining accuracy. Kalign2 is written in C and can also be compiled and called from Python. With some modification, it is possible to communicate with its functions directly from Python, without temporary files.

This allows the greatest efficiency and greatest integration into a Python process. SeqAn and Kalign2 were evaluated against MAFFT, the existing algorithm in use by Du Novo. The aligners were tested by performing a multiple sequence alignment on a duplex read family extracted from a duplex experiment sequencing the whole human mitochondrial genome (Stoler et al. 2016). The family contained 74 reads, 41 single nucleotide substitutions relative to the consensus, and no indels. The number of reads in the alignment was varied from 1 to 74, and the time taken to perform the alignment was measured. Fig. 3 shows the results of this experiment. SeqAn was the slowest at all alignment sizes, with the worst performance at handling of large alignments. It took 58× more time than MAFFT at 10 reads, and 427× more at 40 reads. The fastest for all sizes was Kalign2. At 10 reads, it took less than 10 milliseconds. At 30 reads it was 9× faster, but at 60 reads it was only 4× faster than MAFFT. Since the median family size for an ideal duplex experiment is only around a dozen reads, Kalign2’s advantage is significant and we chose it as the default alignment engine for Du Novo.

**Figure 3.**
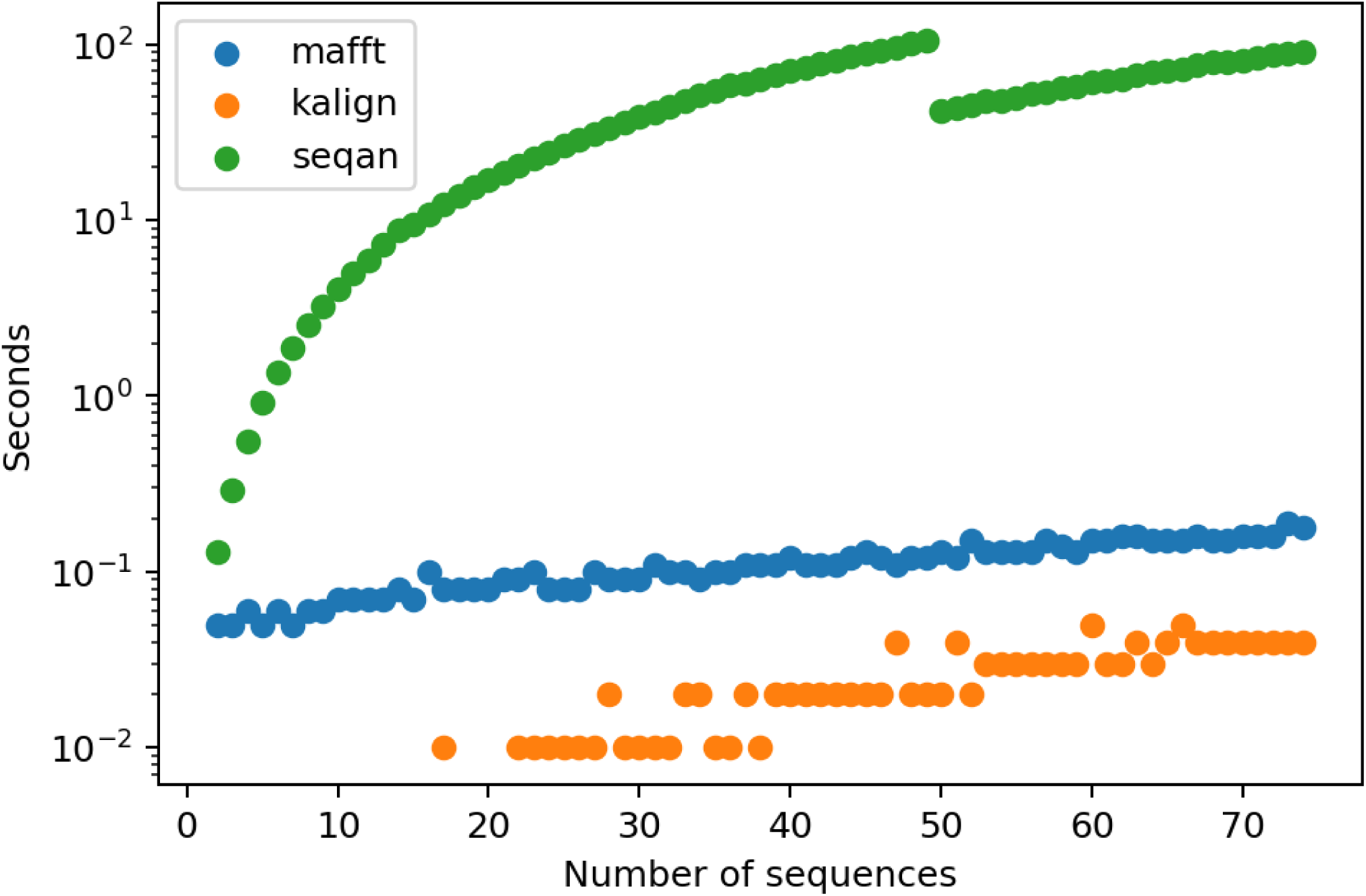
Alignment engine comparison. MAFFT, kalign, and seqan were ran

### Smarter parallelization improves speed

Du Novo uses the multiprocessing Python module for parallel processing. In order to maintain the ordering of the aligned families, the old algorithm would start *N* alignment jobs in parallel, then check each job in order for results. This created a bottleneck at the slowest job: the *N* jobs would take as long as the slowest one. The new algorithm maintains a queue of jobs executing or waiting to be executed. In order to maintain ordering the algorithm keeps an ordered list of submitted jobs. It fills the queue, then begins processing outputs in order of submission. This still requires waiting for all jobs in a batch to finish before continuing, but to reduce the bottleneck, the queue is made larger than the number of available workers. As soon as a job finishes and a worker is freed, it begins work on a new job. This lets new jobs run on CPUs freed by the fastest jobs while the slowest job is still running. If *W* is the number of workers and *M* is a multiplier such that *M*×*W* is the queue size, then a single batch of *M*×*W* jobs will take less time than *M* batches of *W* jobs. Diminishing returns occur as *M* grows, so *M* is set to eight by default. To show the combined effect of the change in alignment and queueing algorithms, Du Novo 2.15, using Kalign2 and the default queue size was compared with Du Novo 0.4, using MAFFT and the old queueing algorithm. Table 2 shows that the combination of the two changes results in an over 9× faster performance at low levels of parallelization. The trend in memory usage is the same as when comparing Kalign2 vs. MAFFT.

**Table 2.** Time and memory usage of different versions of `align-families. py`, using different multiple sequence alignment algorithms. At low levels of parallelization, Kalign2 made the process over 8 times faster, with a memory usage less than twice as much as MAFFT. The new algorithm sped up the tool between 1 and 2.05x. Naturally, at higher levels of parallelization, the reduction of the job queue bottleneck made more of a difference. Memory usage appeared to not be affected, which is expected due to the small size of the job queue compared with the rest of memory usage. To attempt to disentangle the effects of the job queueing algorithm from all the other changes between 0.4 and 2.15, the two versions were compared with all parameters set as similarly as possible. In both cases, the number of`--processes` was set to 32 and MAFFT was used as the aligner. Crucially, the `--queue-size` for the 2.15 version was set to be 32, the same as the number of `--processes`. This approximates the bottleneck in the pre-2.0 version of Du Novo’s job queueing algorithm. Comparing the median of 3 trials of each, the wallclock time of 2.15 was 27% higher than that of 0.4. This could be because of the higher overhead in the more complicated parallelization algorithm, or other changes between 0.4 and 2.15.

| version | aligner | time/<br>memory | CPUs |  |  |  |  |  |
| --- | --- | --- | --- | --- | --- | --- | --- | --- |
|  |  |  | 1 | 2 | 4 | 8 | 16 | 32 |
| 0.4 | MAFFT | time<br>(seconds) | 28,638 | 15,769 | 8,912 | 5,173 | 3,038 | 1,747 |
| 2.15 | MAFFT |  | 28,754 | 14,282 | 7,079 | 3,463 | 1,686 | 854 |
|  | Kalign2 |  | 4,731 | 1,777 | 945 | 600 | 381 | 246 |
| 0.4 | MAFFT | memory<br>(MB) | 23,704 | 12,299 | 6,622 | 3,755 | 2,284 | 1,602 |
| 2.15 | MAFFT |  | 23,927 | 12,599 | 6,850 | 3,985 | 2,541 | 1,810 |
|  | Kalign2 |  | 24,648 | 23,220 | 12,408 | 6,668 | 3,781 | 2,327 |

Next we used the simulated dataset to test whether the change in alignment algorithm affects the accuracy of the pipeline. The simulated experiment was the same, but with 40,000 fragments generated instead of 400,000. Because the input was one homogenous sequence with no minor variants, any differences from the input must be due to incorrect consensus base calls. Using the previous multiple sequence aligner, MAFFT, resulted in an error rate of 0.00563 differences per output base (Table 3). Using Kalign2 instead resulted in 0.00561 differences. Adding barcode error correction improved this figure slightly to 0.00525 while also increasing the yield. The standard pipeline published by Loeb *et al*. (Schmitt et al. 2012) was also compared, resulting in 0.0114 differences per output base.

**Table 3.** Effect of aligner on “correctness”.

| **Method** | **Aligner** | **Barcode error correction** | **Errors per base** |
| --- | --- | --- | --- |
| Du Novo | MAFFT | Uncorrected | 0.563% |
| Du Novo | Kalign2 | Uncorrected | 0.561% |
| Du Novo | Kalign2 | Corrected | 0.525% |
| Loeb | N/A | Uncorrected | 1.140% |

### Using Du Novo

By combining tag error correction with the new alignment engine and parallelization improvements we sought to develop software that can be readily employed by a wide audience of users. To achieve this goal we are distributing Du Novo in three complementary ways:

- Interactive pipeline at http://usegalaxy.org. Here users can upload datasets of any size and process using the complete Du Novo pipeline to produce SSCS and DCS sequences. The Galaxy system contains all tools for downstream processing including mapping and variant calling. To help users effectively use our system we have developed an detailed step-by-step tutorial that can be found here: http://bit.ly/dunovo-tutorial.
- Bioconda package. Du Novo code relies on a number of software components that need to be installed before the tool can be used. Conda package eliminated the need to install these dependencies by automatically installing all components using **conda install** dunovo command (see http://bit.ly/dunovo-bioconda).
- Source code for the package can be found in GitHub at https://github.com/galaxyproject/dunovo. It is distributed under Academic Free License.

## Methods

### Barcode error analysis

The Hamming distance quantifies similarity or dissimilarity between two DNA sequences of equal length by calculating the number of differences between them:

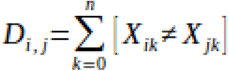

D_*i*,*j*_ is the number of sites where X_*i*_ match and X_*j*_. do not match, *k* is the index of the respective site out of a total number of sites *n*. The input data was in tabular format organized into family size, the sequence of the tag, and the direction of the tag in the SSCS (*ab* = forward or *ba* = reverse). Each tag represents a family of paired-end sequences forming SSCS. Since the whole dataset contained more than one million tags, the comparison of all tags was computationally too demanding. Thus, we parallelized the algorithms and only selected 1,000 random tags from the data set and compared them to the whole dataset to estimate the minimum Hamming distance between tags. A sample of 1,000 tags gives a very similar estimate than the Hamming distance estimated for a sample of 10,000 and ~130,000 tags (Fig. S1). Note that unless noted otherwise, the reported Hamming distance is the minimum Hamming distance of a tag to the other tags in the dataset.

### Error correction

The script `baralign.sh` performs an alignment of all barcodes against themselves (all scripts mentioned in this section can be found at https://github.com/galaxyproject/dunovo). First, it extracts all unique barcode sequences (concatenations of *a*+*b* tags) as FASTA sequences. Then it indexes them, along with their reversed (*b*+*a*) versions, with bowtie-build and aligns them to the index with bowtie `-v 3 --best -a`. This alignment is then read by `correct.py`, which uses the networkx module to constructs graphs where each vertex is a barcode and each edge is a high-quality alignment between two barcodes. The definition of a high-quality alignment can be based on the MAPQ mapping quality, the edit distance given by the NM tag, or distance between the aligned starting positions of the two barcodes. The default values for these filters is 20, 1, and 2, respectively. Then, for each graph, a “correct” barcode is chosen by one of two methods. The default method is to choose the barcode which appears in the largest number of reads. An alternative is to choose the barcode with the most edges to other barcodes.

### Generating simulated duplex data

To validate effectiveness of our approach we have first applied it to a simulated duplex sequencing dataset generated with a duplex sequencing simulator developed to test the correctness of the Du Novo algorithms against known duplex sequencing behavior and sources of errors. It simulates an entire duplex sequencing experiment but taking a reference genome sequence as input, randomly fragmenting it, adding random barcodes to the ends of these fragments, simulating PCR and sequencing errors to produce a set of simulated duplex reads. To randomly fragment the reference sequence, it uses `wgsim` (https://github.com/lh3/wgsim) in error-free mode (options `-e 0 -r 0 -d 0`), using `-1` to set the length of the fragments. Then it simulates random oligomer synthesis to produce duplex barcodes using a uniform 25% probability for each base. It concatenates these oligomers, along with a constant linker sequence, with the fragment sequence to produce starting fragments. These simulated tagged fragments than undergo *in silico* PCR in order to introduce amplification errors. First, a family size is chosen from an empirical distribution observed in a previous duplex experiment. Then, the phylogenetic tree relating these reads is generated. For a family size of *n*, the process starts with *n* reads at the root node representing the original fragment molecule. Each read is randomly assigned to a daughter molecule with 50% probability. Then the process repeats with each daughter, using the number of reads assigned to the daughter instead of *n*. Because amplification efficiencies decline as PCR cycles continue, the probability of replication starts at 1 and is divided by 1.05 each cycle, a realistic value compared with observed reactions (Larionov et al. 2005). Once a tree is generated, errors are simulated at each node and propagated to their descendents. Then, sequencing is simulated, also with errors, and reads are output. A log of the errors is also saved, in order to allow checking results against the “truth”. Unless noted otherwise, simulated data presented here were generated with a sequencing and PCR polymerase error rate of 0.001 errors per base. 25 cycles of PCR were simulated, the fragment lengths were set to 400bp, and the read lengths to 100bp. Using this approach we have generated a datasets containing 400,000 simulated duplex reads and applied our error correction strategy.

### Du Novo 2.0

The basic algorithms in Du Novo 2.15 remain as described in Stoler *et al*. 2016, except the addition of barcode error correction, the Kalign2 multiple sequence aligner, and the replacement of the parallel job queueing algorithm. In all experiments described here, the threshold required to form a consensus base `(make-consensi.py’s --cons-thres)` was 0.7, 3 reads were required to create a consensus sequence `(--min-reads)`, and a PHRED score of at least 25 was required to count a base toward the consensus `(--quai)`.

When consensus reads were filtered, the script `trimmer.py` was used from the `bfx` directory of Du Novo’s distribution. Unless noted otherwise, the script was set to remove the 5’ end of reads when the proportion of N’s in a 10 base window exceeded 0.3 `(--filt-base N --window 10 --thres 0.3)`. If either of the reads in a pair was trimmed to less than 75 bases, both were removed `(--min-length 75)`.

## Acknowledgments

This study has been funded by the funds provided by the Eberly College of Science at the Pennsylvania State University and NIH Grants U41 HG006620 and R01 AI134384-01 as well as NSF ABI Grant 1661497. Funding for R.S., M.H., G.P., and I.T-B was provided by the Linz Institute of Technology (LIT213201001) and the Austrian Science Fund (FWFP30867000).

---

**Figure S1.**
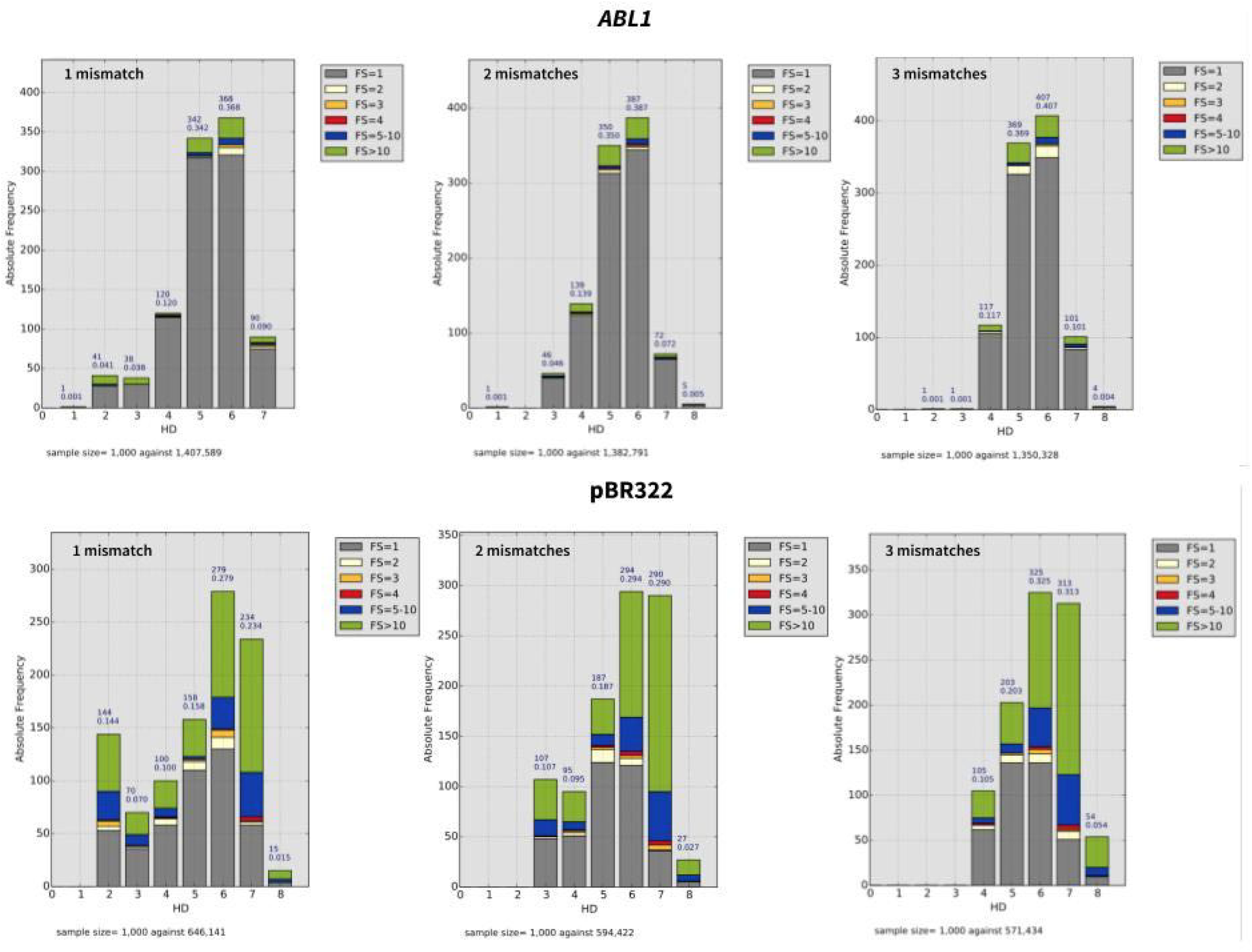
Tag error correction allowing no, 1, 2, or three mismatches. **A**. Shown is the analysis of the sequence differences in the tags (derived from the tabular sequencing file of a targeted genomic region) stratified by family size (FS). The hamming distance (HD) is the number of different nucleotides estimated for a subset of tags (*n*=1000) randomly selected from the overall data (~1 million families) and compared with all the tags in the library. The HD recorded is the smallest HD (minimal HD) of the tag to the rest of the database. **B**. The same data is plotted by family size and stratified by HD. Note that the total number of families decreases as singletons are reassigned to pre-existing families.

